# The ChiS-family DNA-binding domain contains a cryptic helix-turn-helix variant

**DOI:** 10.1101/2020.11.18.389122

**Authors:** Catherine A. Klancher, George Minasov, Ram Podicheti, Douglas B. Rusch, Triana N. Dalia, Karla J. F. Satchell, Matthew B. Neiditch, Ankur B. Dalia

## Abstract

Sequence specific DNA-binding domains (DBDs) are conserved in all domains of life. These proteins carry out a variety of cellular functions, and there are a number of distinct structural domains already described that allow for sequence-specific DNA binding, including the ubiquitous helix-turn-helix (HTH) domain. In the facultative pathogen *Vibrio cholerae*, the chitin sensor ChiS is a transcriptional regulator that is critical for the survival of this organism in its marine reservoir. We have recently shown that ChiS contains a cryptic DBD in its C-terminus. This domain is not homologous to any known DBD, but it is a conserved domain present in other bacterial proteins. Here, we present the crystal structure of the ChiS DBD at a resolution of 1.28 Å. We find that the ChiS DBD contains an HTH domain that is structurally similar to those found in other DNA binding proteins, like the LacI repressor. However, one striking difference observed in the ChiS DBD is that the canonical tight “turn” of the HTH is replaced with an insertion containing a β-sheet, a variant which we term the “helix-sheet-helix”. Through systematic mutagenesis of all positively charged residues within the ChiS DBD, we show that residues within and proximal to the ChiS helix-sheet-helix are critical for DNA binding. Finally, through phylogenetic analyses we show that the ChiS DBD is found in diverse Proteobacterial proteins that exhibit distinct domain architectures. Together, these results suggest that the structure described here represents the prototypical member of the ChiS-family of DBDs.

**Importance:** Regulating gene expression is essential in all domains of life. This process is commonly facilitated by the activity of DNA-binding transcription factors. There are diverse structural domains that allow proteins to bind to specific DNA sequences. The structural basis underlying how some proteins bind to DNA, however, remains unclear. Previously, we showed that in the major human pathogen *Vibrio cholerae* the transcription factor ChiS directly regulates gene expression through a cryptic DNA binding domain. This domain lacked homology to any known DNA-binding protein. In the current study, we determined the structure of the ChiS DNA binding domain (DBD) and find that the ChiS-family DBD is a cryptic variant of the ubiquitous helix-turn-helix (HTH) domain. We further demonstrate that this domain is conserved in diverse proteins that may represent a novel group of transcriptional regulators.

## Introduction

The intestinal pathogen *Vibrio cholerae* natively resides in the aquatic environment and can cause disease if ingested in the form of contaminated food or drinking water. In the aquatic environment, *V. cholerae* commonly associates with the chitinous surfaces of crustacean zooplankton (1). Chitin is an abundant source of carbon and nitrogen for marine bacteria, including *V. cholerae* (2, 3). In addition, chitin serves as a cue to induce horizontal gene transfer by natural transformation in this species (4). Thus, *Vibrio*-chitin interactions are critical for this facultative pathogen to thrive and evolve in its environmental reservoir.

Chitin is sensed in *V. cholerae* by the hybrid histidine kinase ChiS (5–7). In response to chitin, ChiS activates the expression of the chitin utilization program. This regulon includes the *chb* operon, which is required for the uptake and degradation of the chitin disaccharide chitobiose. In a recent study, we showed that unlike most histidine kinases, ChiS is capable of directly binding to DNA to regulate the expression of the *chb* operon (5). This finding was particularly surprising because ChiS is not predicted to encode a DNA-binding domain *via* primary sequence homology (BLAST (8)) or structural predictions (Phyre2 (9)). In the current study, we sought to understand the structural basis for ChiS DNA binding. To that end, we determined the structure of the ChiS DBD and found that it encodes a distinct variant of the canonical helix-turn-helix domain, which we term a “helix-sheet-helix”.

## Results and Discussion

### The C-terminus of ChiS (ChiS^1024-1129^) is sufficient to bind P_chb_

Previous work from our group demonstrates that ChiS is a noncanonical hybrid histidine kinase that contains a DBD at its C-terminus (**Fig. 1A**) (5). In that study, we found that the C-terminal 106 amino acids of ChiS (ChiS^1024-1129^) were necessary and sufficient to bind to the *chb* promoter *in vivo. We* further showed that ChiS binds directly to two binding sites within the *chb* operon promoter (P_*chb*_) to activate the expression of this locus. To confirm that ChiS^1024-1129^ was sufficient to bind DNA, we purified this domain and tested DNA-binding activity *in vitro* by electrophoretic mobility shift assays (EMSAs). We found that ChiS^1024-1129^ bound to a wild-type P_*chb*_ promoter probe, but not to a probe in which the two ChiS binding sites were mutated, suggesting that this domain is sufficient to bind to DNA in a sequencespecific manner (**Fig. 1B** and **Fig. S1**). Thus, based on our *in vivo* and *in vitro* analysis, we refer to ChiS^1024-1129^ as the ChiS DBD.

**Figure 1.**
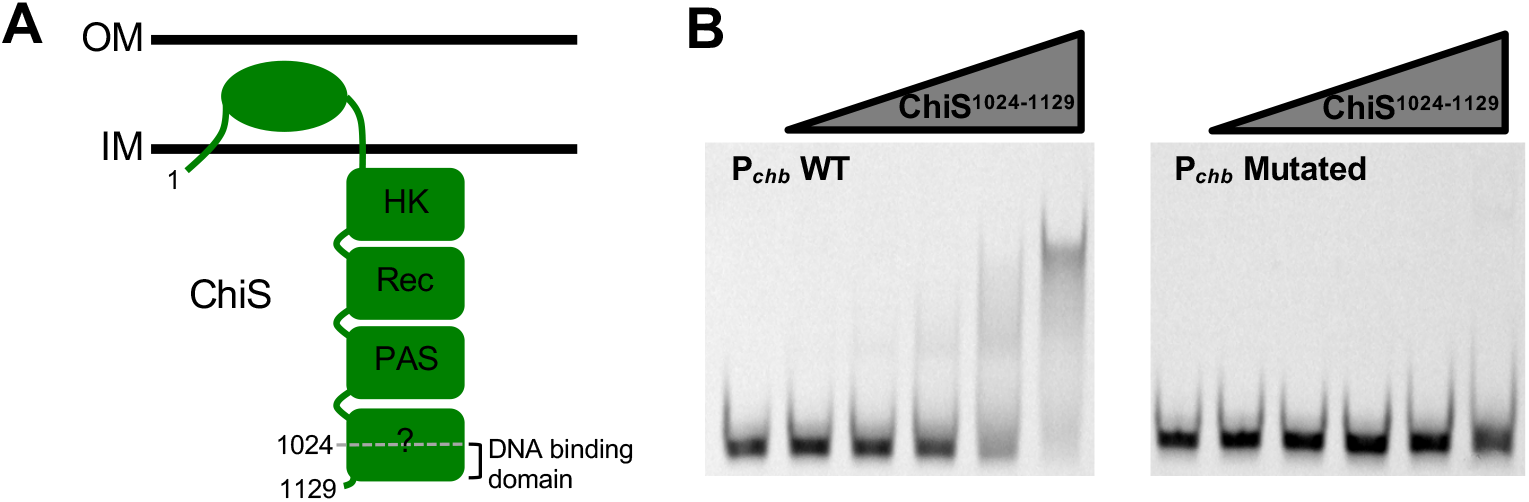
The C-terminus of ChiS (ChiS^1024-1129^) is sufficient to bind P_chb_. **A**) Diagram of the domain architecture for the hybrid histidine kinase ChiS. ChiS contains a histidine kinase (HK) domain, a receiver domain (Rec), a PAS domain, and a domain that does not have homology to known domains. Residues 1024-1129 were previously shown to be sufficient to bind P_*chb*_ *in vivo* (5). **B**) A fragment of the ChiS C-terminus (ChiS^1024-1 129^) was purified and assessed for DNA binding activity by EMSA. Purified protein was incubated with the indicated Cy5-labeled 60 bp probes containing sequence from P_*chb*_ encompassing the two ChiS binding sites (CBSs). The probe sequence wad WT (P_*chb*_ WT) or the CBSs were both mutated (P_*chb*_ Mutated). **See Figure S1A** for a promoter map and the probe sequences used. The concentration of ChiS used (from left to right) was 0 nM, 25 nM, 50 nM, 1θ0 nM, 200 nM, and 400 nM. Data are representative of two independent experiments.

### Identification of positively charged residues in the ChiS DBD that are critical for DNA binding and transcriptional activation of P_chb_

As mentioned above, ChiS is not predicted to encode a DNA-binding domain. This is based on *in silico* searches using the ChiS DBD. With BLAST, no conserved domains were detected in the ChiS DBD (8). Further, Phyre2 predicted structural models were of very low confidence, and none of the hits identified contained a known DNA-binding domain (9).

To characterize interactions between the ChiS DBD and DNA, we first tried to identify residues important for DNA binding. The positively charged amino acids arginine (R) and lysine (K) commonly interact with the negatively charged DNA backbone and can also make critical contacts with nucleotide bases (10). Thus, we mutated every R and K residue in the ChiS DBD to a glutamine (Q), to ablate their charge but maintain, to a reasonable extent, the steric properties of the side group. To determine how these mutations affected ChiS activity, we introduced them into full-length FLAG-tagged ChiS (5), and assessed the ability of each mutant to bind to DNA *in vivo* (by chromatin immunoprecipitation, or ChIP) and to activate P_*chb*_ expression (using a P_*chb*_-GFP reporter). Full-length ChiS was used for these experiments because this construct is functional for both DNA-binding and transcriptional activation of P_*chb*_, whereas the ChiS DBD is only functional for DNA-binding (5). We found that all mutations to ChiS reduced P_*chb*_-GFP activation to varying degrees (**Fig. 2**). Most mutants were able to facilitate partial activation of P_*chb*_ and, correspondingly, partially enriched for P_*chb*_ by ChIP, indicating that they were binding to the promoter *in vivo*. Some mutants (R1068Q, R1074Q, K1078Q, R1090Q, and R1092Q) did not bind to P_*chb*_ DNA *in vivo* and resulted in complete loss of P_*chb*_ expression. All mutants still produced ChiS protein as assessed by Western blot analysis (**Fig. S2**), however, we cannot exclude the possibility that these single amino acid substitutions result in protein misfolding. Collectively, these data identify a subset of positively charged residues in the ChiS DBD that are likely critical for DNA binding and subsequent transcriptional activation of the *chb* operon.

**Figure 2.**
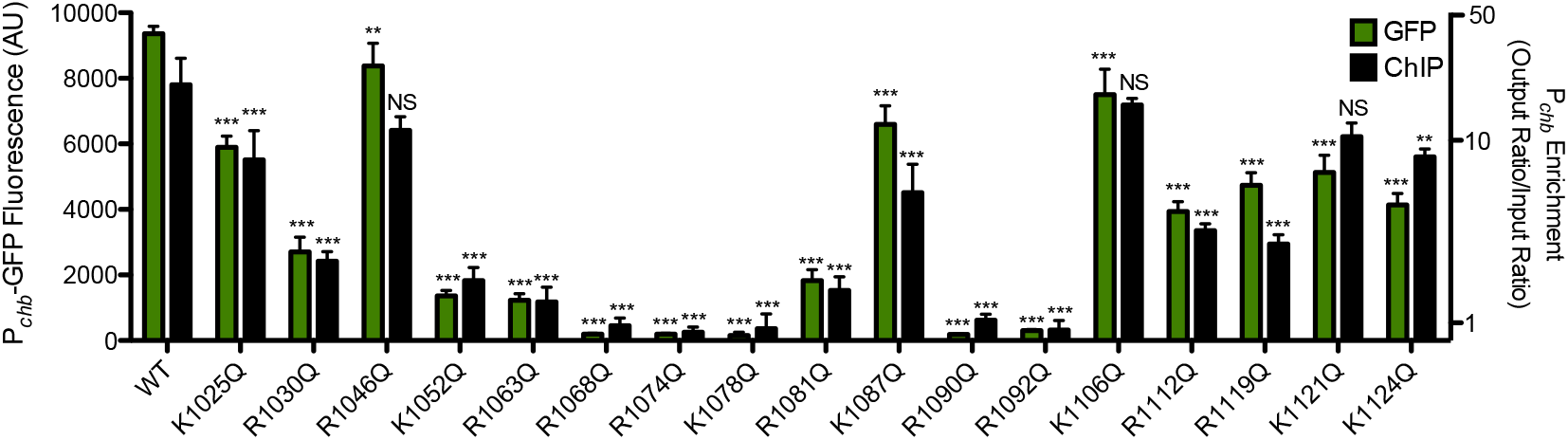
Identification of positively charged residues in the ChiS DBD that are critical for DNA binding and transcriptional activation of P_chb_. All lysines and arginines in the ChiS DNA binding domain were individually mutated to a glutamine and ChiS was assessed for (1) transcriptional activation of a P_*chb*_-GFP reporter (green bars; left Y-axis) and (2) ChiS binding to P_*chb*_ *in vivo* by chromatin immunoprecipitation (ChIP) (black bars; right Y-axis). ChiS can be activated with its native inducer, chitin, or by deletion of its periplasmic regulator, CBP; here, ChiS was activated artificially by deleting CBP. Data are the result of at least three independent biological replicates and are shown as the mean ± SD. Statistical markers indicated directly above bars indicate comparisons to the WT. Statistical comparisons were made by one-way ANOVA with Tukey’s post test. ***, p < 0.001; ** p < 0.01; NS, not significant.

### Structure of the ChiS DNA binding domain reveals a variant of the helix-turn-helix

We next sought to determine the structure of the ChiS DBD to further explore how ChiS interacts with DNA. Since no structures for close sequence homologs were available in the Protein Data Bank (PDB) to serve as search models for molecular replacement, we used the Single-wavelength Anomalous Dispersion (SAD) technique to determine initial phases. Selenomethionine was used as the replacement for methionine. Anomalous data were collected from a single crystal (**Tables S1 and S2**). The crystal diffracted to 1.28 Å resolution and belonged to the orthogonal C222_1_ space group with unit cell parameters of a=51.91Å, b=78.61Å, c=72.37Å, α=β=γ=90.00°. There was one polypeptide chain in the asymmetric unit. The structure includes 105 out of 106 residues of the protein (1024 – 1128), two uncleavable residues of the purification tag, four sulfate ions (SO_4_^2−^), one 2-(2-hydroxyethyloxy)ethanol molecule (PEG), two formic acids molecules (FMT) and 200 water molecules (HOH). Only the C-terminal E1129 was disordered in the structure and was not included in the final model.

The structure of the ChiS DBD revealed that it contains a fold that is reminiscent of the canonical helix-turn-helix (HTH) used by diverse DNA-binding proteins (**Fig. 3A-B**). The basic HTH domain consists of a trihelical bundle where the second and third helix encompass the namesake “helixturn-helix” (11). The two helices that compose the HTH are connected *via* a relatively short linker that forms a sharp turn, which is a characteristic feature of this domain. Helix 3 from the HTH is generally inserted into the major groove of DNA, forming the principal DNA-protein interface. Alignment of the trihelical bundle from ChiS with the DNA-bound structure of the LacI repressor (PDB: 1EFA (12); RMSD of modeled C_α_ carbons = 3.514) revealed a similar spatial arrangement for each helix (**Fig. 3C**). In addition, LacI and ChiS have similar electrostatic properties, suggesting that a positively charged protein interface that interacts with DNA is a conserved feature of both proteins (**Fig. S3**). Notably, however, the ChiS HTH has an insertion containing two anti-parallel β-strands connected by a turn between helix 2 and helix 3 that form a β-sheet (**Fig. 3B-D**). Structural insertion between these helices is not typical; thus, the sheet feature found here is a distinct variant of the HTH which we refer to as a “helix-sheet-helix”. Comparison of the ChiS trihelical bundle to other structures in the PDB using the DALI server (13) did not reveal any structures that resemble the helix-sheet-helix described here, suggesting that this structure represents a new variant of the HTH.

**Figure 3.**
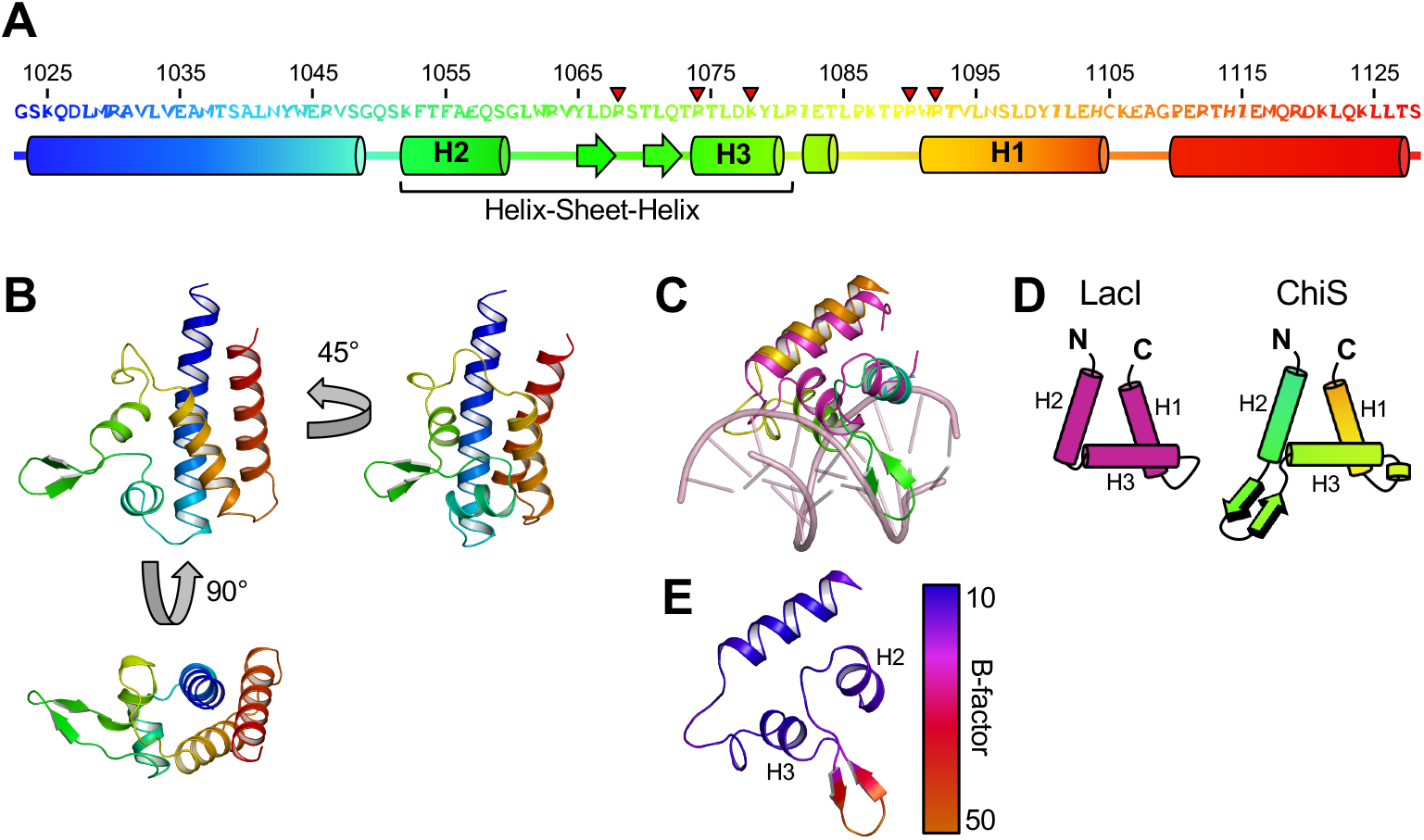
Structure of the ChiS DNA binding domain reveals a variant of the helix-turn-helix. **A**) Domain architecture of the ChiS DNA binding domain. The primary sequence of the ChiS DBD is shown. Helices are depicted as cylinders, while sheets are depicted as arrows. The five R and K residues found to be critical for DNA-binding are denoted by red arrowheads. **B**) Crystal structure of the ChiS DNA binding domain. The structural elements are color-coded as depicted in the primary sequence in **A**. **C**) Alignment of the ChiS trihelical bundle (rainbow) with the LacI triehlical bundle bound to the LacI operator site (PDB: 1EFA; pink). Alignment of alpha carbons gave an RMSD of 3.514. **D**) Cartoon representations of the trihelical bundle from LacI and ChiS. Helices are labeled with nomenclature presented in Aravind, *et. al*. (11). **E**) Structure of the ChiS trihelical bundle colored to represent B-factor. Helices found in the helix-sheet-helix motif (H2 and H3) are indicated.

Alignment of the ChiS DBD to LacI also revealed that the sheet within the ChiS helix-sheet-helix domain runs along the major groove (**Fig. 3C, 4A**), though it sterically conflicts with the DNA bases. This may suggest that the ChiS DBD takes on a slightly different conformation when bound to DNA. Consistent with this idea, the β-sheet insertion has the highest B-factor (a measure of structural motion) in the ChiS DBD structure, indicating that it is relatively flexible (**Fig. 3E**). Despite the elevated B-factor in this region, an omit map indicates that the antiparallel beta strands of the sheet are strongly supported by the data collected (**Fig. S4**). We speculate that this β-sheet is stabilized in the major groove when the ChiS DBD is bound to DNA. The unique helix-sheet-helix feature of the ChiS C-terminal domain may also explain why it was not previously identified as a DBD by structure prediction algorithms like Phyre2.

**Figure 4.**
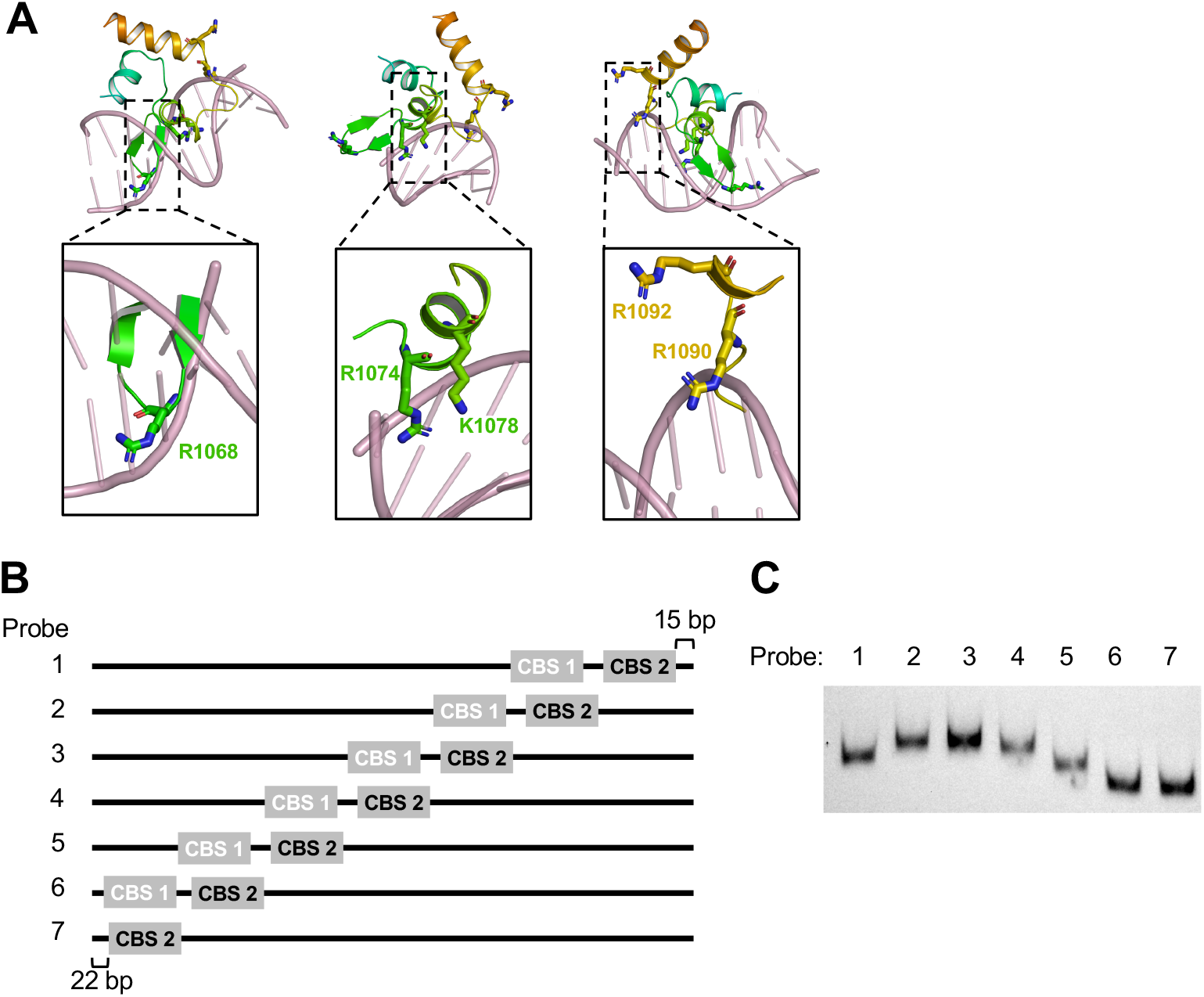
ChiS may bind to intrinsically bent DNA. **A**) Model of the ChiS trihelical bundle bound to double-stranded DNA from the alignment shown in **Figure 3C**. Side chains for the residues critical for DNA binding (R1068, R1074, K1078, R1090, and R1092) are shown and indicated. **B**) Diagram of the 7 (distinct 230 bp probes used in **C**. ChiS binding site 1 (CBS 1) was mutated (white text) and ChiS binding site 2 (CBS 2) was left intact (black text). CBS 2 was shifted by 30 bp between each probe. **C**) The DNA probesdiagtammed in **B** were labeled with Cy5 and separated by native PAGE: in the absence of ChiS protein.

### ChiS may bind to intrinsically bent DNA

Above, we identified five residues (R1068, R1074, K1078, R1090, and R1092) that were critical for the ChiS DBD to bind to DNA. Mapping these residues onto the ChiS DBD structure revealed that all five residues were found within the trihelical bundle that forms the helix-sheet-helix (**Fig. 4A**), which is consistent with this domain playing a critical role in DNA binding. Specifically, these residues were located in the β-sheet of the helix-sheet-helix (R1068), helix 3 (R1074, K1078), and helix 1 (R1090, R1092).

Most residues critical for DNA binding activity (R1068, R1074, K1078, R1090) were in close proximity to DNA on our modeled alignment. Based on the model, we can speculate on the DNA contacts made by these residues. R1068 is found in the sheet of the helix-sheet-helix, which, as stated above, sterically conflicts with DNA bases on our modeled alignment. Thus, it is unclear whether R1068 would make contact with the DNA backbone or with the nucleotide bases. R1074 and K1078 model closest to the nucleotide bases, suggesting that these residues may be critical for base pair recognition. R1090 on the other hand potentially makes contacts with the DNA backbone.

While the above residues modeled closely to DNA, one residue (R1092), was distant from the DNA (**Fig. 4A**). Many transcription factors bend DNA upon binding to their target site (14, 15). Thus, one possible explanation for the critical role of R1092 is that the P_*chb*_ promoter is bent when bound by ChiS, which would allow for R1092 to come into close contact with DNA. To test this idea, we carried out a classic *in vitro* gel mobility shift assay to test DNA bending (16). This assay operates on the basis that the location of a bend within a DNA molecule alters its mobility during native PAGE analysis (17, 18). DNA probes that contain a bend in the middle of the probe exhibit the lowest mobility, while probes with the bend closer to one end show the highest mobility. Thus, we designed 7 DNA probes of equal length that gradually shifted the position of the ChiS binding sites within the *chb* promoter (**Fig. 4B** and **Fig. S1**). First, we ran these probes in the absence of ChiS protein and found that they ran at different mobilities where the probes with the ChiS binding sites in the middle exhibited the lowest mobility (**Fig. 4C**). This suggested that the *chb* promoter likely has an intrinsic bend that is centered around the ChiS binding sites. The mobility pattern observed for these DNA probes did not change when incubated with the purified ChiS DBD (**Fig. S5**), suggesting that binding of the DNA probe by ChiS does not further bend the promoter. We propose that the *chb* promoter has an intrinsic bend, which may allow residues in the ChiS DBD, like R1092, to directly interact with DNA. The intrinsic bend found in the *chb* promoter may increase the affinity of ChiS for this region of DNA; indeed, DNA bending has been shown to increase the affinity of certain transcription factors for their DNA binding site (19).

### The ChiS family DNA binding domain is associated with variable domain arrangements in diverse proteins

Above, we show that the ChiS DBD represents a cryptic variant of an HTH domain. As noted previously, the ChiS DBD is found in proteins other than homologs of ChiS (5). To more fully catalog proteins that contain this domain, we generated a profile Hidden Markov Model (HMM) to the ChiS DBD and screened for its presence among eubacterial genomes. A profile HMM is a position-specific scoring system that can effectively encode the variation in a training set of representative peptide sequences, and then find similar sequences from a much larger and more distantly related dataset compared to tools that do not require training, such as BLAST (20, 21).

This analysis revealed that the ChiS DBD is present in diverse Proteobacterial genomes (**Spreadsheet S1**). The vast majority of hits from our search were direct homologs of ChiS (3242/3829 = 84.7%), however, many proteins exhibited distinct domain architectures (587/3829 = 15.3%) (**Fig. 5A**). Strikingly, the ChiS DBD was found exclusively at the C-terminus in all of these proteins and was commonly associated with sensory domains (**Fig. 5A**). The residues found to be critical for DNA binding in **Figure 2** had varying degrees of conservation with ChiS family DBD containing proteins (**Fig. 5B**). R1068 is poorly conserved, suggesting that this residue may be involved in sequence specific interactions with DNA. Consistent with this idea, R1068 may interact closely with the nucleotide bases (**Figure 3A**). R1074 and R1092 are somewhat conserved, and K1078 and R0190 are very well conserved across several proteins. This suggests that these residues of the ChiS-family DBD may be required for general DNA interactions and do not contribute to sequence specificity. In general, the helix-sheet-helix is highly conserved across these diverse proteins (**Fig. 5B, Spreadsheet S1**), and even the most dissimilar ChiS DBD homolog (MAC43155.1, bit score of 43.5; 22.6% identical, 43.4% similar to the ChiS DBD) still threaded (9) remarkably well onto the trihelical bundle of the ChiS DBD structure (RMSD of modeled C_α_ carbons = 0.002) (**Fig. S6**). Thus, we suggest that ChiS is the founding member for a new group of DNA-binding transcription factors whose activity are regulated by diverse sensory inputs.

**Figure 5.**
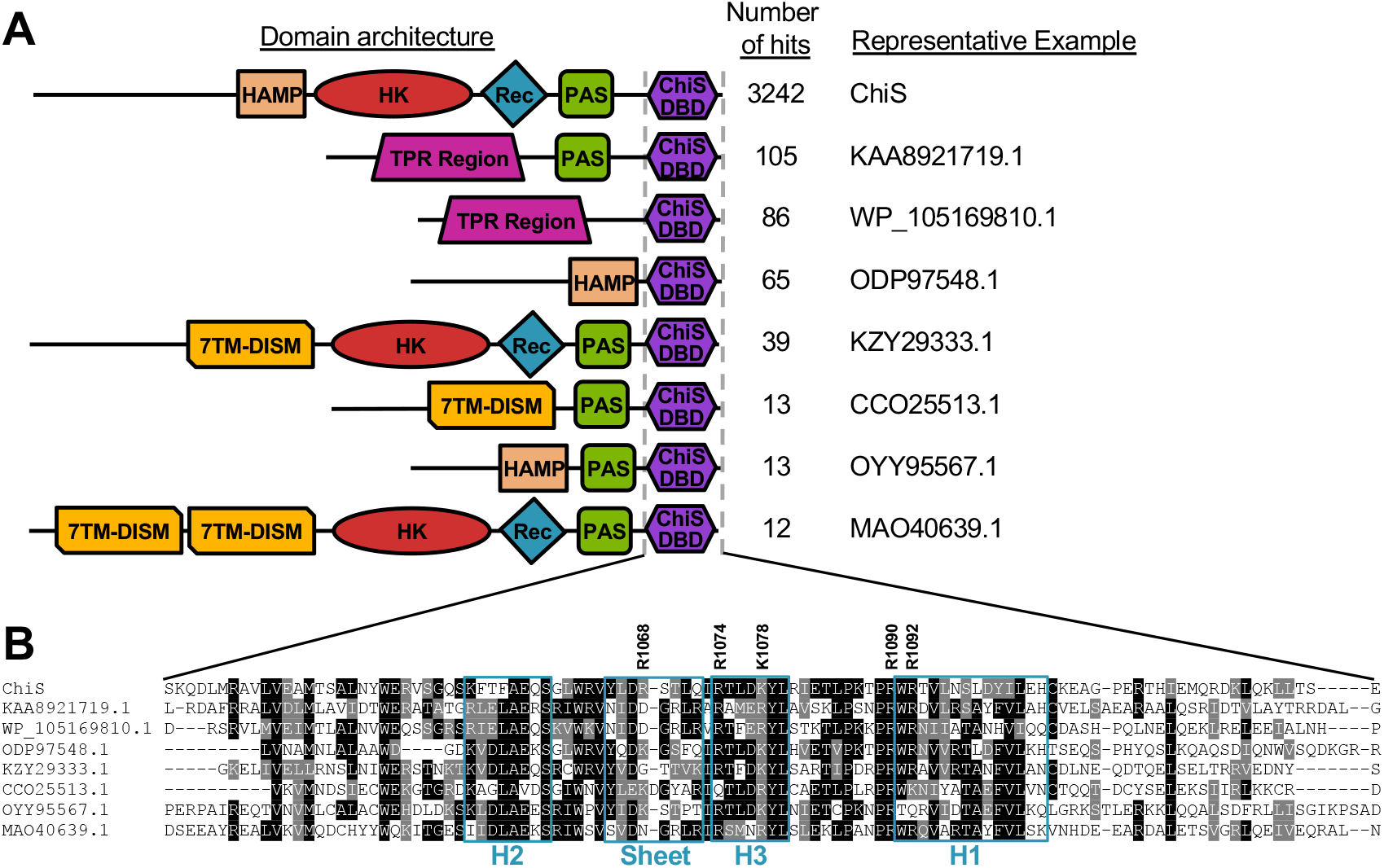
The ChiS family DBD is found in diverse proteins with distinct domain architectures among Proteobacterial genomes. **A**) Diagrams of the most abundant protein architectures containing the ChiS family DBD. Protein domains shown are HAMP, Histidine Kinase (HK), Receiver (Rec), Per-Arnt-Sim (PAS), Tetratricopeptide Repeat (TPR), 7 Transmembrane Receptors with Diverse Intracellular Signaling Modules (7TMR-DISM), and the ChiS family DNA binding domain (ChiS DBD). For a complete list of hits containing the indicated architectures see **Spreadsheet S1**. **B**) Alignment of the primary sequences of the ChiS family DBD in the indicated proteins are shown. Residues in black are identical, while those in gray are similar. The sequence for helix 1 (H1), helix 2 (H2), helix 3 (H3), and the sheet of the helix-sheet-helix are boxed in teal. The five Ft and K residues found to be critical for DNA-binding in ChiS are also indicated.

In this study, we have characterized the first member of the ChiS-family of DBDs. Though many DBDs have been extensively studied, this work demonstrates that subtle structural variants of canonical DBDs can be difficult to identify by structural prediction algorithms, like Phyre2. Further, the findings here suggest that many proteins with DBDs may currently be eluding detection. As many residues required for DNA-binding in the ChiS DBD are well-conserved, our data suggest that there is a common mechanism of binding DNA within the ChiS-family DBDs. Our work also indicates that the canonical tight turn of the HTH is not a critical feature for sequence-specific DNA-binding, and further highlights the diversity in structural solutions that can allow for this type of activity. While we have generated a putative model of the ChiS DBD bound to DNA in this study, it remains unclear how the sheet within the helix-sheet-helix contributes to sequence-specific DNA binding. Solving the structure of a ChiS-family DBD bound to DNA will be the focus of future work.

## Materials & Methods

### Bacterial strains and culture conditions

All *V. cholerae* strains used in this study are derived from the El Tor strain E7946 (22). *V. cholerae* strains were grown in LB medium and on LB agar supplemented when necessary with carbenicillin (20 μg/mL), kanamycin (50 μg/mL), spectinomycin (200 μg/mL), and/or trimethoprim (10 μg/mL). See **Table S3** for a detailed list of mutant strains used in this study.

### Generating mutant strains

*V. cholerae* mutant constructs were generated using splicing-by-overlap extension exactly as previously described (23). See **Table S4** for all of the primers used to generate mutant constructs in this study. Mutant *V. cholerae* strains were generated by chitin-dependent natural transformation and cotransformation exactly as previously described (24). Mutant strains were confirmed by PCR and/or sequencing.

### Cloning, protein production and purification

The *chiS^1024-1129^* construct was cloned into an Amp^R^ pET15b-based vector using the FastCloning method (25). This vector appended a TEV cleavable 6x His tag onto the N-terminus of ChiS^1024-1129^. Vector and inserts were amplified using the primers listed in **Table S4**. The plasmid was transformed into *E. coli* BL21(DE3) (Magic) cells (26) and the protein was expressed in M9 media (High Yield M9 Se-Met media, Medicilon Inc.). The starting overnight culture was grown in LB medium supplemented with 130 μg/mL ampicillin and 50 μg/mL kanamycin at 37°C and 220 rpm. The next day, M9 medium supplemented with 200 μg/mL ampicillin and 50 μg/mL kanamycin was inoculated with the overnight culture (1:100 dilution) and incubated at 37°C and 220 rpm. Protein expression was induced at OD_600_=1.8-2.0 by the addition of 0.5 mM isopropyl β-d-1-thiogalactopyranoside and the culture was further incubated at 25°C, 200 rpm for 14 hours (27). The cells were harvested by centrifugation at 6,000 xg for 10 minutes, resuspended to 0.2 g/mL in lysis buffer (50 mM Tris pH 8.3, 0.5 M NaCl, 10% glycerol, 0.1% IGEPAL CA-630) and frozen at −30°C until purification.

Frozen pellets were thawed and sonicated at 50% amplitude, in a 5s on, 10s off cycle for 20 min at 4°C. The lysate was clarified by centrifugation at 18,000 xg for 40 minutes at 4°C and the supernatant was collected. The protein was purified in one step by IMAC followed by size exclusion chromatography using ÅKTAxpress system (GE Healthcare) as previously described with some modifications (28). The cell extract was loaded into a His-Trap FF (Ni-NTA) column with loading buffer (10 mM Tris-HCl pH 8.3, 500 mM NaCl, 1 mM Tris (2-carboxyethyl) phosphine (TCEP), 5% glycerol) and the column was washed with 10 column volumes of loading buffer and 10 column volumes of washing buffer (10 mM Tris-HCl pH 8.3, 1 M NaCl, 25 mM imidazole, 5% glycerol). Protein was eluted with elution buffer (10 mM Tris pH 8.3, 500 mM NaCl, 1 M imidazole), loaded onto a Superdex 200 26/600 column, separated in loading buffer, collected, and analyzed by PAGE. The 6x His tag was cleaved with recombinant TEV protease in a ratio of 1:20 (protein:protease) overnight at room temperature. The cleaved protein was separated from uncleaved protein, recombinant TEV protease, and 6x His tag peptide by Ni-NTA-affinity chromatography using loading buffer followed by loading buffer with 25 mM imidazole. The cleaved protein was collected in the flow-through in both the loading buffer and the loading buffer with 25 mM imidazole. Both fractions were analyzed by PAGE for 6x His tag cleavage, concentrated to 6-8 mg/mL, and set up for crystallization.

### Crystallization, data collection, structure solution and refinement

The protein from both fractions (collected in flow through and in 25 mM imidazole) was set up at 6-8 mg/mL in loading buffer containing 0 or 500 mM NaCl as 2 μL crystallization drops (1 μL protein: 1 μL reservoir solution) in 96-well plates (Corning) using commercial Classics II, PACT and JCSG+ (QIAGEN) crystallization screens. Diffraction quality crystal of the protein collected with 25 mM imidazole grown from the condition with 0.2 M lithium sulfate, 0.1 M Bis-Tris, pH 5.5, 25%(w/v) PEG 3350 (Classics II, #74) was flash frozen in liquid nitrogen for data collection.

The crystals were screened, and data were collected at the Life Sciences-Collaborative Access Team (LS-CAT) beamline F at the Advanced Photon Source (APS) of the Argonne National Laboratory. A total of 300 diffraction images were indexed, integrated and scaled using HKL-3000 (29). The structure was determined with the HKL3000 structure solution package using anomalous signal from selenomethionine (Se-Met). The initial model went through several rounds of refinement in REFMAC v. 5.8.0258 (30) and manual corrections in Coot (31). The water molecules were generated using ARP/wARP (32) and ligands were added to the model manually during visual inspection in Coot. Translation–Libration–Screw (TLS) groups were created by the TLSMD server (33) and TLS corrections were applied during the final stages of refinement. MolProbity (34) was used for monitoring the quality of the model during refinement and for the final validation of the structure. The structure was deposited to the Protein Data Bank (https://www.rcsb.org/) with the assigned PDB code 7KPO. Structural diagrams were drawn from PDB files using the PyMOL Molecular Graphics System v2.4 (Schrödinger, Inc).

### Electrophoretic mobility shift assay (EMSA)

Binding reactions contained 10 mM Tris HCl pH 7.5, 1 mM EDTA, 10 mM KCl, 1 mM DTT, 50 μg/mL BSA, 0.1 mg/mL salmon sperm DNA, 5% glycerol, 1 nM of a Cy5 labeled DNA probe, and purified ChiS DBD at the indicated concentrations (diluted in 10 mM Tris pH 7.5, 10 mM KCl, 1 mM DTT, and 5% glycerol). Reactions were incubated at room temperature for 20 minutes in the dark, then electrophoretically separated on polyacrylamide gels in 0.5x Tris Borate EDTA (TBE) buffer at 4°C. Gels were imaged for Cy5 fluorescence on a Typhoon-9210 instrument. Cy5-labeled P_*chb*_ probes were made by Phusion PCR, where Cy5-dCTP was included in the reaction at a level that would result in incorporation of 1–2 Cy5 labeled nucleotides in the final probe as previously described (23).

### Measuring GFP reporter fluorescence

GFP fluorescence was determined essentially as previously described (35). Briefly, single colonies were picked and grown in LB broth at 30°C for 18 hours. Cells were then washed and resuspended to an OD_600_ of 1.0 in instant ocean medium (7 g/L; Aquarium Systems). Then, fluorescence was determined using a BioTek H1M plate reader with excitation set to 500 nm and emission set to 540 nm.

### Chromatin immunoprecipitation (ChIP)-qPCR assays

ChIP assays were carried out exactly as previously described (5). Briefly, overnight cultures were diluted to an OD_600_ of 0.08 and then grown for 6 hours at 30°C. Cultures were crosslinked using 1% paraformaldehyde, then quenched with a 1.2 molar excess of Tris. Cells were washed with PBS and stored at −80°C overnight. The next day, cells were resuspended in lysis buffer (1 x FastBreak cell lysis reagent (Promega), 50 μg/mL lysozyme, 1% Triton X-100, 1 mM PMSF, and 1x protease inhibitor cocktail; 100x inhibitor cocktail contained the following: 0.07 mg/mL phosphoramidon (Santa Cruz), 0.006 mg/mL bestatin (MPbiomedicals/Fisher Scientific), 1.67 mg/mL AEBSF (DOT Scientific), 0.07 mg/mL pepstatin A (Gold Bio), 0.07 mg/mL E64 (Gold Bio)) and then lysed by sonication, resulting in a DNA shear size of ~500 bp. Lysates were incubated with Anti-FLAG M2 Magnetic Beads (Sigma), washed to remove unbound proteins, and then bound protein-DNA complexes were eluted off with SDS. Samples were digested with Proteinase K, then crosslinks were reversed. DNA samples were cleaned up and used as template for quantitative PCR (qPCR) using iTaq Universal SYBR Green Supermix (Bio-Rad) and primers specific for the genes indicated (see **Table S4** for primers) on a Step-One qPCR system. Standard curves of genomic DNA were included in each experiment and were used to determine the abundance of each amplicon in the input (derived from the lysate prior to ChIP) and output (derived from the samples after ChIP). Primers to amplify *rpoB* served as a baseline control in this assay because ChiS does not bind this locus. Data are reported as ‘Fold Enrichment’, which is defined as the ratio of P_*chb*_ / *rpoB* found in the output divided by the same ratio found in the input.

### Western blot analysis

Strains were grown as described for ChIP assays, pelleted, resuspended, and boiled in 1x SDS PAGE sample buffer (110 mM Tris pH 6.8, 12.5% glycerol, 0.6% SDS, 0.01% Bromophenol Blue, and 2.5% β-mercaptoethanol). Proteins were separated by SDS polyacrylamide gel electrophoresis, then transferred to a PVDF membrane, and probed with rabbit polyconal α-FLAG (Sigma) or mouse monoclonal α-RpoA (Biolegend) primary antibodies. Blots were then incubated with α-rabbit or α-mouse HRP conjugated secondary antibodies, developed using Pierce ECL 529 Western Blotting Substrate (ThermoFisher), and imaged on a ProteinSimple Fluorchem E instrument.

### Bioinformatic identification of eubacterial proteins with putative ChiS DBD domains

The DBD sequence segments from the protein sequences of seven ChiS DNA binding domain homologs (THB81618.1, OGG93021.1, OUR95018.1, WP_084205767.1, ODU31202.1, WP_070993003.1, WP_078715702.1) (5) were aligned using MUSCLE version 3.8.31 (36). The resulting multiple sequence alignment was turned into a profile HMM which was searched against the eubacterial subset (taxonomy id: 2) of NCBI non-redundant protein sequence database using HMMER version 3.2.1 (http://hmmer.org/), requiring the alignment length to be at least 90. Among the hits, protein sequences tagged “partial” in their FASTA headers were excluded. Domain architectures for the remaining hits were obtained from the NLM conserved domain database (37). Any protein hits with regions aligned to the DNA binding domain HMM overlapping with known annotated functional domains were excluded. The resulting ChiS DBD homolog protein sequences were clustered using cd-hit ver. v4.8.1-2019-0228 (38) (parameters: -M 0 -g 1 -s 0.8 -c 0.4 -n 2 -d 500). Clusters identified by cd-hit were further grouped together by manually analyzing the domain architecture of hits as shown in **Fig. 5A**. Only clusters containing 10 or more representatives were grouped, while the remaining proteins were left unassigned. For a list of all proteins containing a putative ChiS DBD, see **Spreadsheet S1**.

## Acknowledgements

We thank Dipankar Sen, Julia van Kessel, and Ryan Chaparian for helpful discussions. This work was supported by grant R35GM128674 from the National Institutes of Health (to ABD) and, in part, with Federal funds from the Department of Health and Human Services. National Institutes of Health, National Institute of Allergy and Infectious Diseases under Contract No. HHSN272201700060C. This research used resources of the Advanced Photon Source, a U.S. Department of Energy (DOE) Office of Science User Facility operated for the DOE Office of Science by Argonne National Laboratory under Contract No. DE-AC02-06CH11357. Use of the LS-CAT Sector 21 was supported by the Michigan Economic Development Corporation and the Michigan Technology Tri-Corridor (Grant 085P1000817). This research was supported in part by Lilly Endowment, Inc., through its support for the Indiana University Pervasive Technology Institute.

---

**Figure S1.**
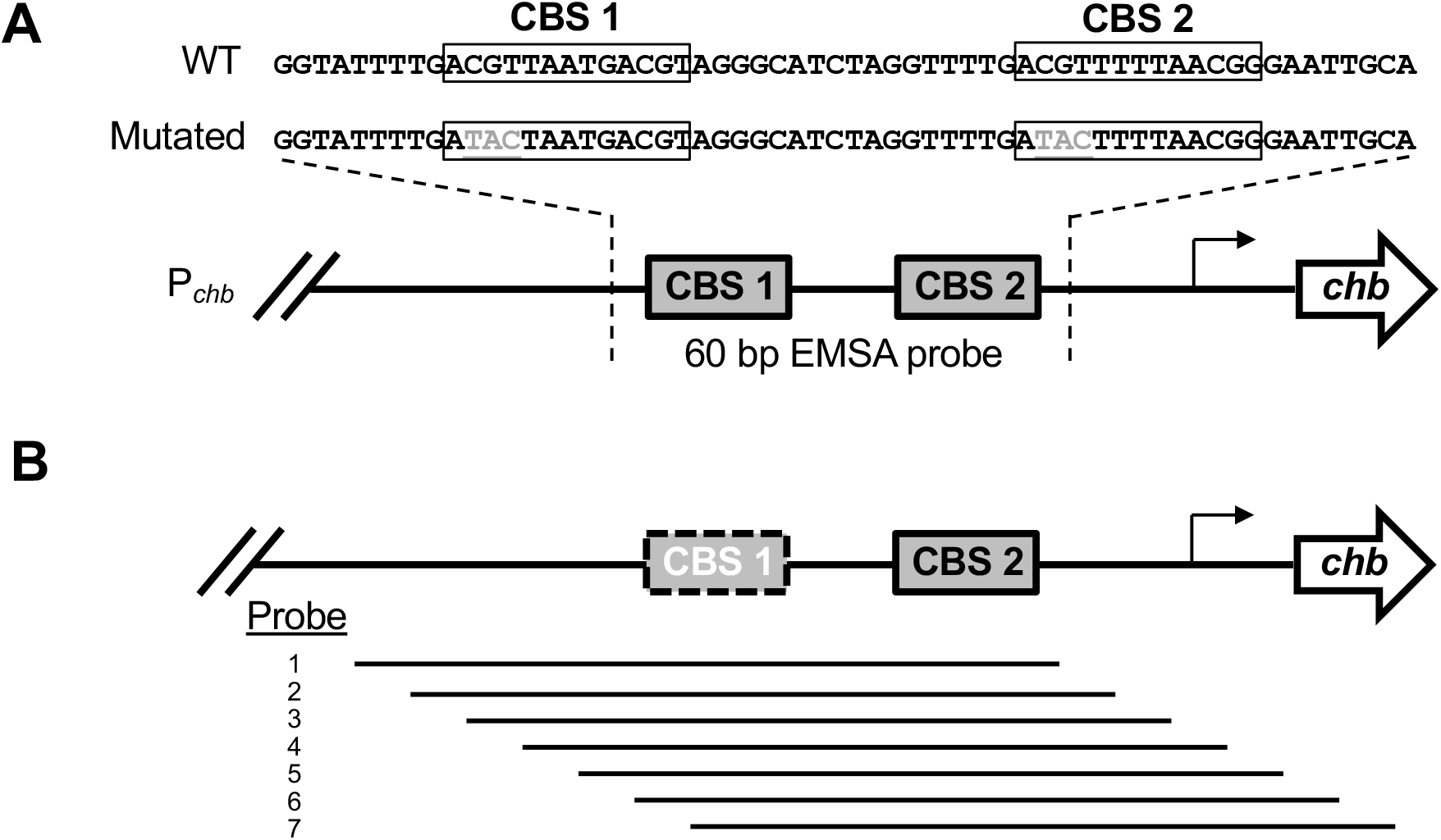
Diagrams of EMSA probes used in this study. **A**) Promoter map of *chb* with the region of P_*chb*_ used for the EMSAs shown in **Figure 1B** indicated. The exact probe sequences are shown above the promoter map. ChiS binding site (CBS) are boxed and the mutations used to disrupt the CBSs are shown in gray and underlined. **B**) Promoter map of *chb* with the region of P_*chb*_ used for the EMSA shown in **Figure 4B** and **Figure S5**. CBS 1 was mutated (white text, dotted line) in all probes used.

**Figure S2.**
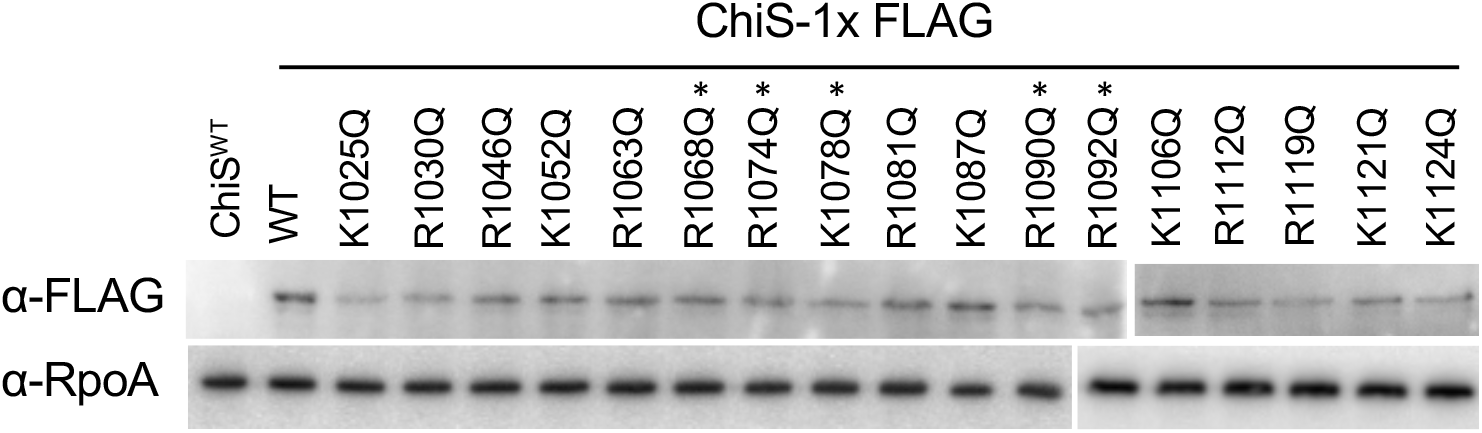
Mutations to the ChiS DNA binding domain does not prevent expression of ChiS. Strains expressing the indicated ChiS-FLAG point mutations were assessed for expression by Western blot with anti-FLAG and anti-RpoA (loading control) antibodies. Asterisks above ChiS point mutants indicate the mutations found to be critical for the DNA binding activity of ChiS as shown in **Figure 2**.

**Figure S3.**
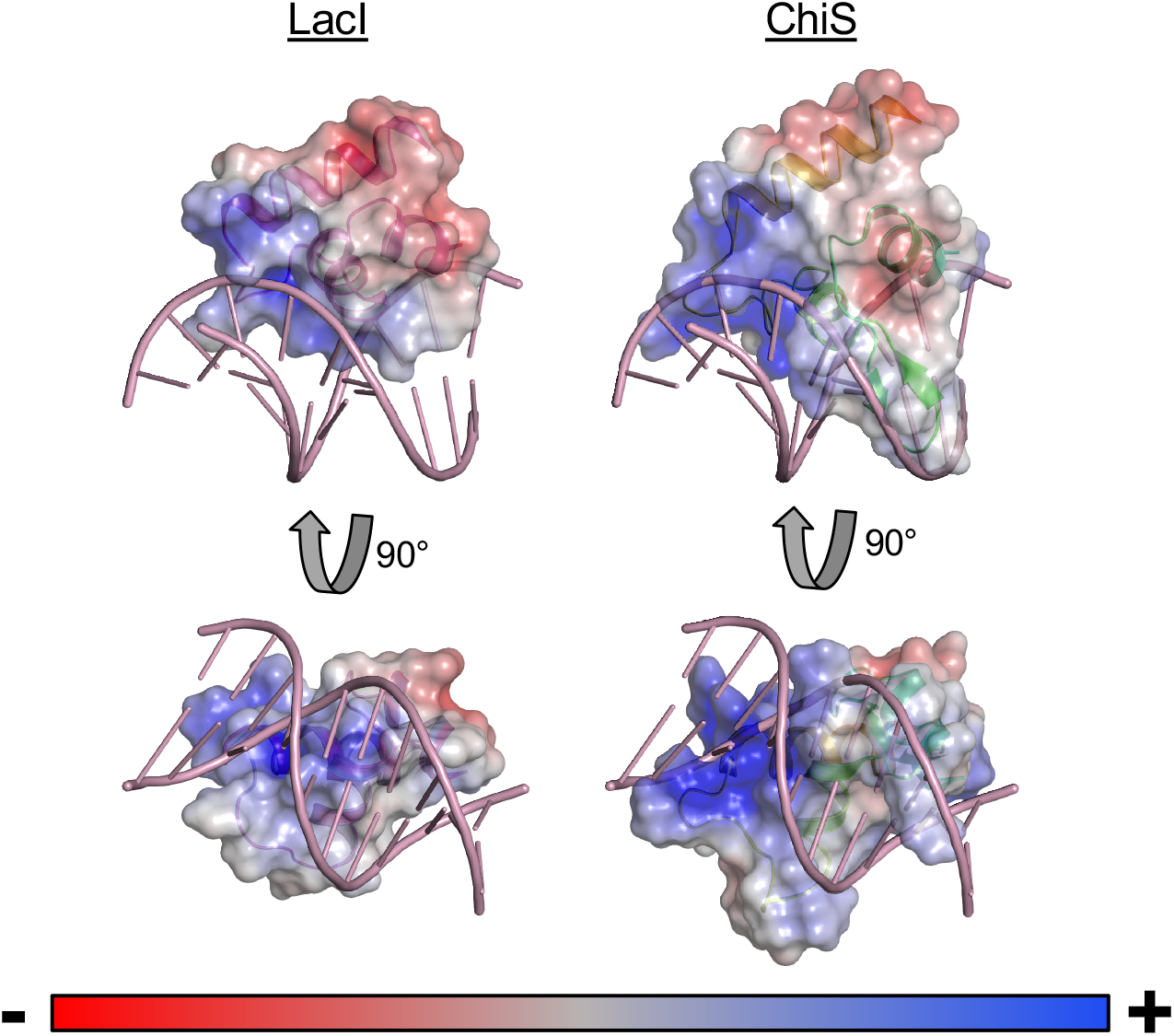
The electrostatic surface pattern of the LacI helix-turn-helix and ChiS helixsheet-helix are similar. Electrostatic maps of the LacI helix-turn-helix DNA-bound structure and the ChiS helix-sheet-helix DNA-bound model. The DNA shown is from the LacI structure, and the ChiS DBD was modeled onto DNA by alignment to LacI as shown in **Figure 3C**. Regions of the protein surface colored in red are negatively charged, while those shown in blue are positively charged.

**Figure S4.**
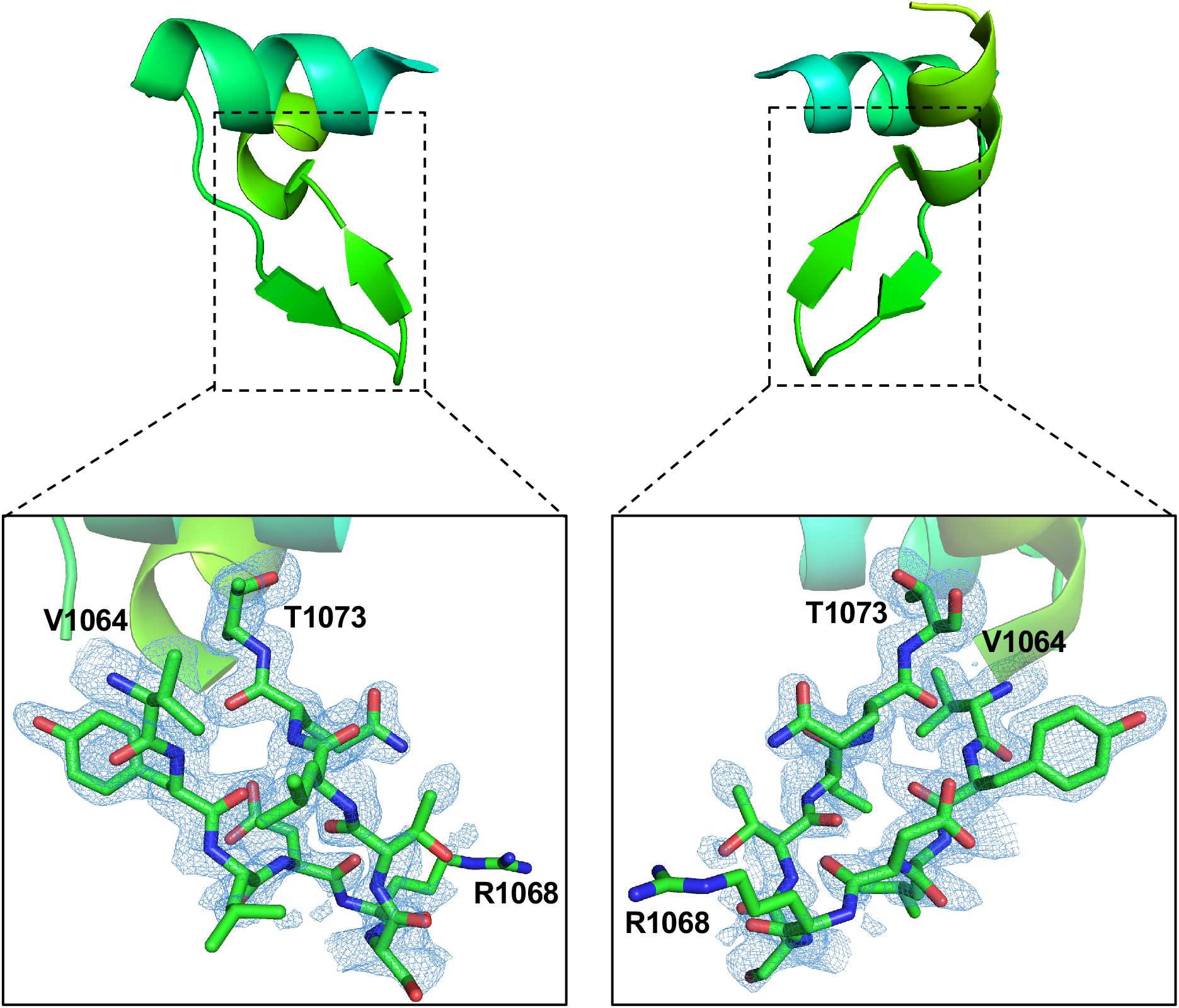
Electron density map of the β-sheet insertion with in the helix-sheet-helix. An omit F_o_ - F_c_ map contoured at the 2.0 sigma level (omitted V1064-T1073) was generated and mapped onto the β-sheet insertion of the ChiS helix-sheet-helix. The map reveals that despite the high B-factor in this region, the structure modeled within the antiparallel beta strands are strongly supported by the data collected. The modeled side chains of the residues in the turn of the beta-turn-beta insertion, however, are less clearly resolved, which may be due to the flexibility of this region.

**Figure S5.**
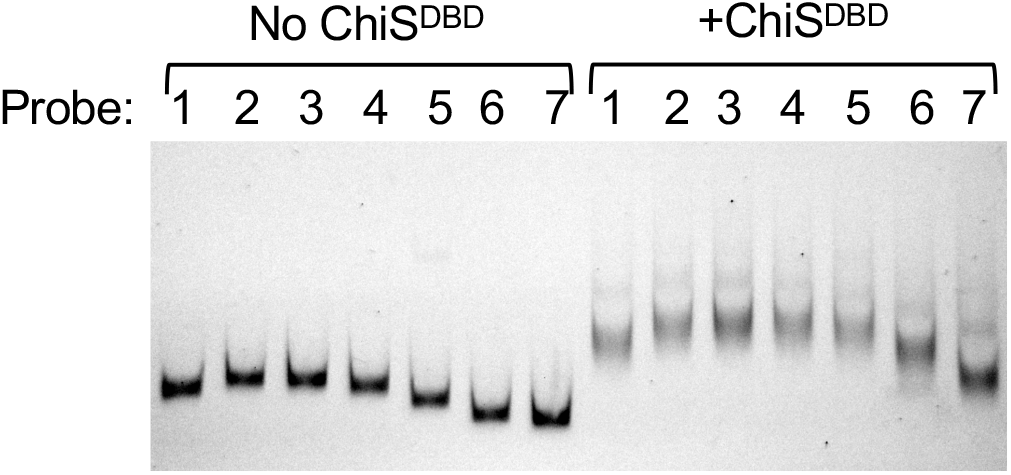
ChiS protein does not further bend the P_chb_ promoter. The probes shown in **Figure S1B** were incubated in the absence (No ChiS^DBD^) or presence (+ChiS^DBD^) of 400 nM ChiS^DBD^.

**Figure S6.**
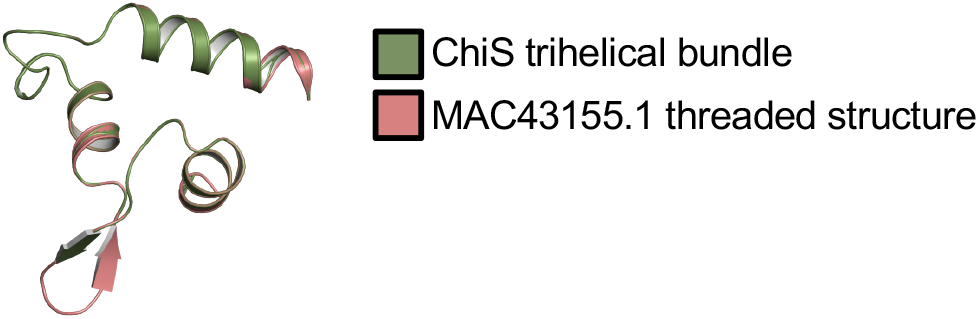
The most dissimilar ChiS DBD homolog threads onto the trihelical bundle of the ChiS DBD structure. The sequence of the ChiS-family DBD from MAC43155.1 was threaded onto the crystal structure of the ChiS DBD using Phyre2 (9). Alignment of alpha carbons gave an RMSD of 0.002.

**Table S1.** Data collection and processing Values in parentheses are for the outer shell.

|  |  |
| --- | --- |
| Diffraction source | Beamline 21ID-F, APS |
| Wavelength (Å) | 0.97872 |
| Temperature (K) | 100 |
| Detector | MAR Mosaic 300 mm CCD |
| Space group | C222<sub>1</sub> |
| a, b, c (Å) | 51.91, 78.61, 72.37 |
| α, β, γ (°) | 90.00, 90.00, 90.00 |
| Resolution range (Å) | 30.00 - 1.28 (1.30 - 1.28) |
| No. of unique reflections | 38,590 (1,866) |
| Completeness (%) | 99.7 (97.3) |
| Multiplicity | 6.9 (4.6) |
| $\langle I/\sigma(I) \rangle$ | 28.3 (2.3) |
| R<sub>r.i.m.</sub><sup>†</sup> | 0.032 (0.386) |
| CC<sub>1/2</sub><sup>††</sup> | (0.637) |
| Overall B factor from Wilson plot (Å<sup>2</sup>) | 14.7 |

<sup>†</sup>Estimated $R_{r.i.m.} = R_{merge}[N/(N - 1)]^{1/2}$ , where N is the data multiplicity.
<sup>††</sup> Pearson's Correlation Coefficient (Karplus & Diederichs, 2012).

**Table S2.** Structure refinement Values in parentheses are for the outer shell.

|  |  |
| --- | --- |
| Resolution range (Å) | 25.97 - 1.28 (1.31 - 1.28) |
| Completeness (%) | 99.7 (98.3) |
| No. of reflections, working set | 36,482 (2,634) |
| No. of reflections, test set | 1,885 (138) |
| Final R<sub>work</sub> | 0.144 (0.214) |
| Final R<sub>free</sub> | 0.180 (0.238) |
| No. of non-H atoms |  |
| Protein | 989 |
| Ligand | 33 |
| Water | 200 |
| Total | 1,222 |
| R.m.s. deviations |  |
| Bonds (Å) | 0.005 |
| Angles (°) | 1.217 |
| Average B factors (Å<sup>2</sup>) |  |
| Protein | 17.8 |
| Ligand | 31.6 |
| Water | 31.3 |
| Ramachandran plot |  |
| Favored regions (%) | 99.0 |
| Additionally allowed (%) | 1.0 |
| Outliers (%) | 0.0 |

**Table S3.**
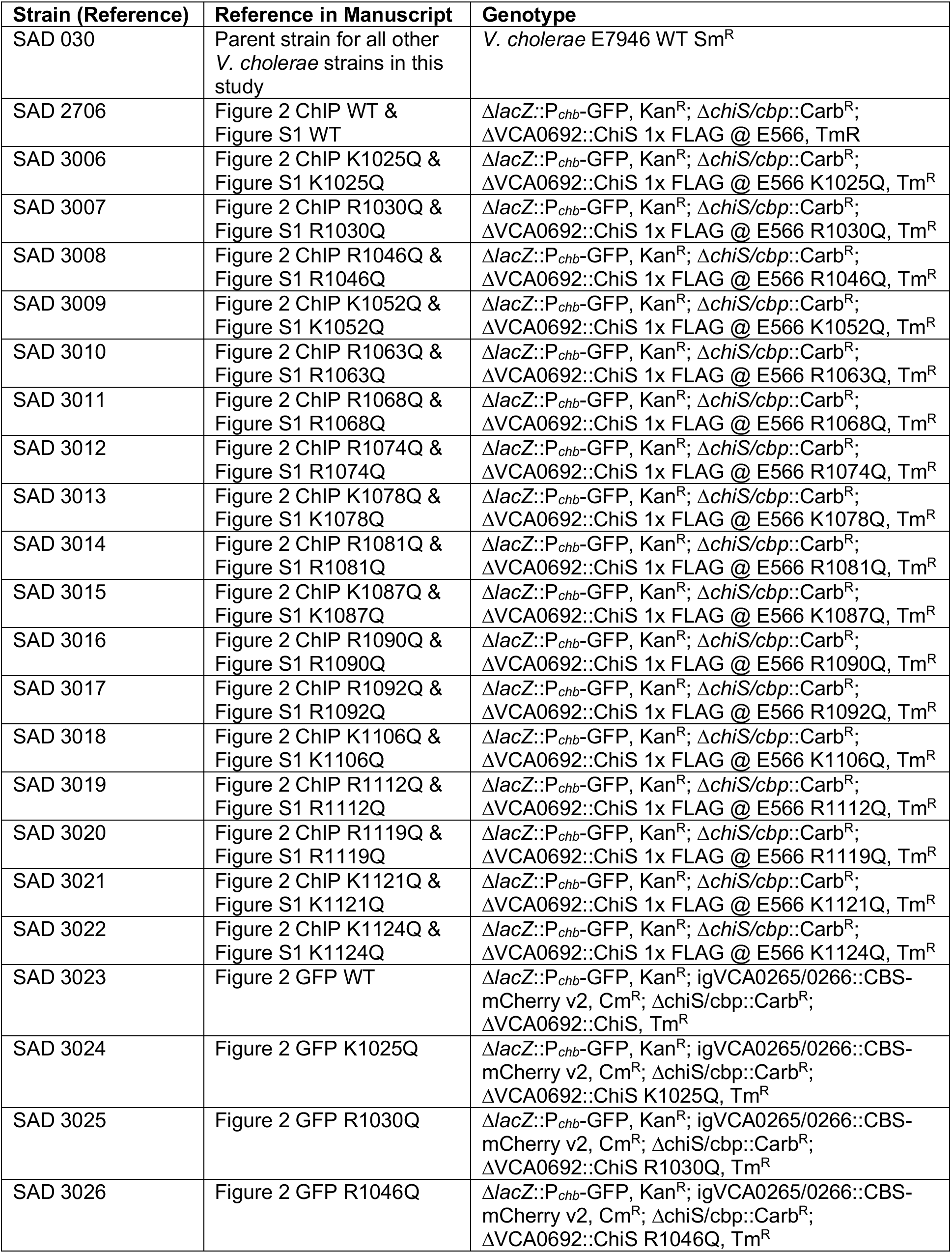
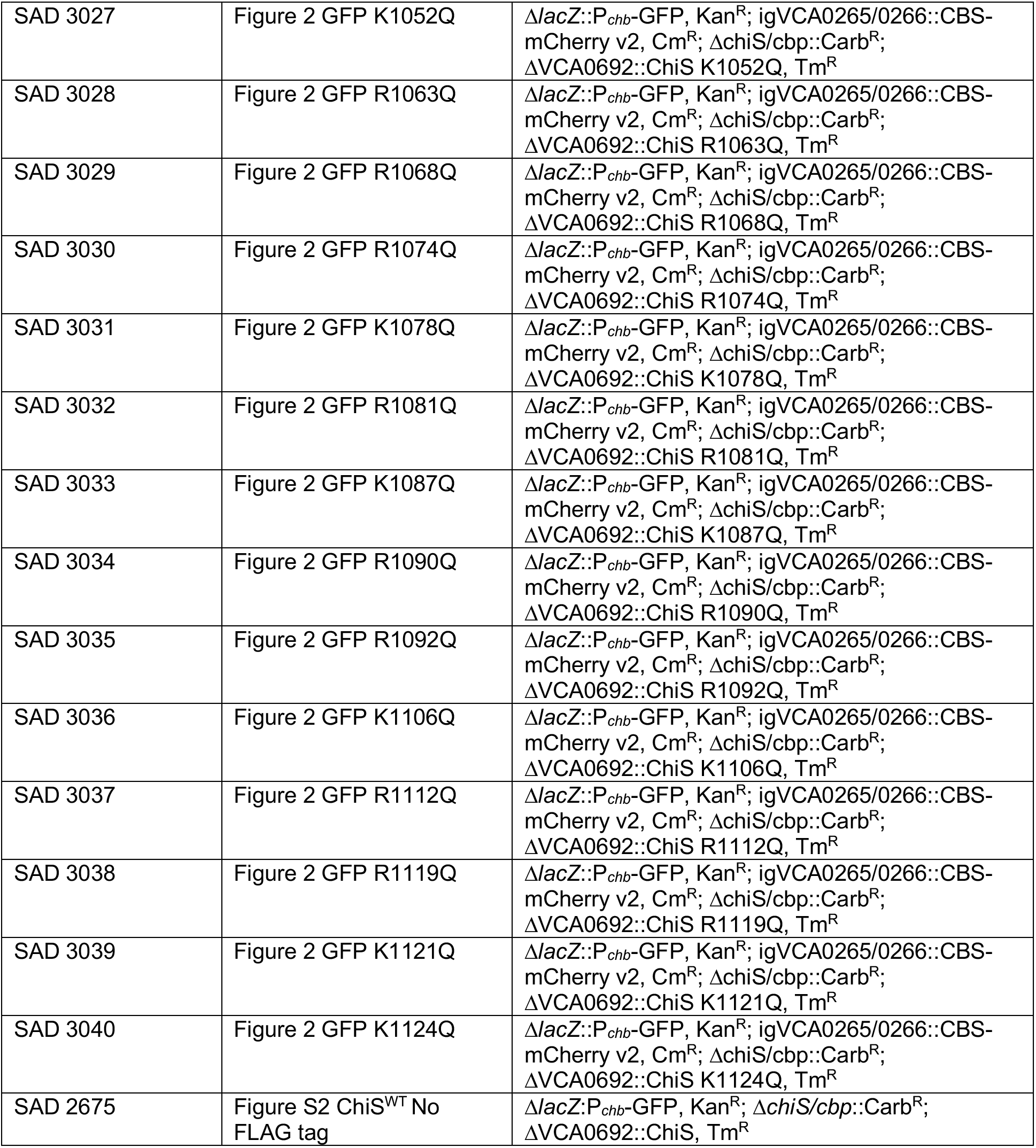
Strains used in this study.

| Strain (Reference) | Reference in Manuscript | Genotype |
| --- | --- | --- |
| SAD 030 | Parent strain for all other *V. cholerae* strains in this study | *V. cholerae* E7946 WT Sm<sup>R</sup> |
| SAD 2706 | Figure 2 ChIP WT & Figure S1 WT | $\Delta lacZ::P_{chb}$-GFP, Kan<sup>R</sup>; $\Delta chiS/cbp::Carb^R$; $\Delta VCA0692::ChiS$ 1x FLAG @ E566, Tm<sup>R</sup> |
| SAD 3006 | Figure 2 ChIP K1025Q & Figure S1 K1025Q | $\Delta lacZ::P_{chb}$-GFP, Kan<sup>R</sup>; $\Delta chiS/cbp::Carb^R$; $\Delta VCA0692::ChiS$ 1x FLAG @ E566 K1025Q, Tm<sup>R</sup> |
| SAD 3007 | Figure 2 ChIP R1030Q & Figure S1 R1030Q | $\Delta lacZ::P_{chb}$-GFP, Kan<sup>R</sup>; $\Delta chiS/cbp::Carb^R$; $\Delta VCA0692::ChiS$ 1x FLAG @ E566 R1030Q, Tm<sup>R</sup> |
| SAD 3008 | Figure 2 ChIP R1046Q & Figure S1 R1046Q | $\Delta lacZ::P_{chb}$-GFP, Kan<sup>R</sup>; $\Delta chiS/cbp::Carb^R$; $\Delta VCA0692::ChiS$ 1x FLAG @ E566 R1046Q, Tm<sup>R</sup> |
| SAD 3009 | Figure 2 ChIP K1052Q & Figure S1 K1052Q | $\Delta lacZ::P_{chb}$-GFP, Kan<sup>R</sup>; $\Delta chiS/cbp::Carb^R$; $\Delta VCA0692::ChiS$ 1x FLAG @ E566 K1052Q, Tm<sup>R</sup> |
| SAD 3010 | Figure 2 ChIP R1063Q & Figure S1 R1063Q | $\Delta lacZ::P_{chb}$-GFP, Kan<sup>R</sup>; $\Delta chiS/cbp::Carb^R$; $\Delta VCA0692::ChiS$ 1x FLAG @ E566 R1063Q, Tm<sup>R</sup> |
| SAD 3011 | Figure 2 ChIP R1068Q & Figure S1 R1068Q | $\Delta lacZ::P_{chb}$-GFP, Kan<sup>R</sup>; $\Delta chiS/cbp::Carb^R$; $\Delta VCA0692::ChiS$ 1x FLAG @ E566 R1068Q, Tm<sup>R</sup> |
| SAD 3012 | Figure 2 ChIP R1074Q & Figure S1 R1074Q | $\Delta lacZ::P_{chb}$-GFP, Kan<sup>R</sup>; $\Delta chiS/cbp::Carb^R$; $\Delta VCA0692::ChiS$ 1x FLAG @ E566 R1074Q, Tm<sup>R</sup> |
| SAD 3013 | Figure 2 ChIP K1078Q & Figure S1 K1078Q | $\Delta lacZ::P_{chb}$-GFP, Kan<sup>R</sup>; $\Delta chiS/cbp::Carb^R$; $\Delta VCA0692::ChiS$ 1x FLAG @ E566 K1078Q, Tm<sup>R</sup> |
| SAD 3014 | Figure 2 ChIP R1081Q & Figure S1 R1081Q | $\Delta lacZ::P_{chb}$-GFP, Kan<sup>R</sup>; $\Delta chiS/cbp::Carb^R$; $\Delta VCA0692::ChiS$ 1x FLAG @ E566 R1081Q, Tm<sup>R</sup> |
| SAD 3015 | Figure 2 ChIP K1087Q & Figure S1 K1087Q | $\Delta lacZ::P_{chb}$-GFP, Kan<sup>R</sup>; $\Delta chiS/cbp::Carb^R$; $\Delta VCA0692::ChiS$ 1x FLAG @ E566 K1087Q, Tm<sup>R</sup> |
| SAD 3016 | Figure 2 ChIP R1090Q & Figure S1 R1090Q | $\Delta lacZ::P_{chb}$-GFP, Kan<sup>R</sup>; $\Delta chiS/cbp::Carb^R$; $\Delta VCA0692::ChiS$ 1x FLAG @ E566 R1090Q, Tm<sup>R</sup> |
| SAD 3017 | Figure 2 ChIP R1092Q & Figure S1 R1092Q | $\Delta lacZ::P_{chb}$-GFP, Kan<sup>R</sup>; $\Delta chiS/cbp::Carb^R$; $\Delta VCA0692::ChiS$ 1x FLAG @ E566 R1092Q, Tm<sup>R</sup> |
| SAD 3018 | Figure 2 ChIP K1106Q & Figure S1 K1106Q | $\Delta lacZ::P_{chb}$-GFP, Kan<sup>R</sup>; $\Delta chiS/cbp::Carb^R$; $\Delta VCA0692::ChiS$ 1x FLAG @ E566 K1106Q, Tm<sup>R</sup> |
| SAD 3019 | Figure 2 ChIP R1112Q & Figure S1 R1112Q | $\Delta lacZ::P_{chb}$-GFP, Kan<sup>R</sup>; $\Delta chiS/cbp::Carb^R$; $\Delta VCA0692::ChiS$ 1x FLAG @ E566 R1112Q, Tm<sup>R</sup> |
| SAD 3020 | Figure 2 ChIP R1119Q & Figure S1 R1119Q | $\Delta lacZ::P_{chb}$-GFP, Kan<sup>R</sup>; $\Delta chiS/cbp::Carb^R$; $\Delta VCA0692::ChiS$ 1x FLAG @ E566 R1119Q, Tm<sup>R</sup> |
| SAD 3021 | Figure 2 ChIP K1121Q & Figure S1 K1121Q | $\Delta lacZ::P_{chb}$-GFP, Kan<sup>R</sup>; $\Delta chiS/cbp::Carb^R$; $\Delta VCA0692::ChiS$ 1x FLAG @ E566 K1121Q, Tm<sup>R</sup> |
| SAD 3022 | Figure 2 ChIP K1124Q & Figure S1 K1124Q | $\Delta lacZ::P_{chb}$-GFP, Kan<sup>R</sup>; $\Delta chiS/cbp::Carb^R$; $\Delta VCA0692::ChiS$ 1x FLAG @ E566 K1124Q, Tm<sup>R</sup> |
| SAD 3023 | Figure 2 GFP WT | $\Delta lacZ::P_{chb}$-GFP, Kan<sup>R</sup>; igVCA0265/0266::CBS-mCherry v2, Cm<sup>R</sup>; $\Delta chiS/cbp::Carb^R$; $\Delta VCA0692::ChiS$, Tm<sup>R</sup> |
| SAD 3024 | Figure 2 GFP K1025Q | $\Delta lacZ::P_{chb}$-GFP, Kan<sup>R</sup>; igVCA0265/0266::CBS-mCherry v2, Cm<sup>R</sup>; $\Delta chiS/cbp::Carb^R$; $\Delta VCA0692::ChiS$ K1025Q, Tm<sup>R</sup> |
| SAD 3025 | Figure 2 GFP R1030Q | $\Delta lacZ::P_{chb}$-GFP, Kan<sup>R</sup>; igVCA0265/0266::CBS-mCherry v2, Cm<sup>R</sup>; $\Delta chiS/cbp::Carb^R$; $\Delta VCA0692::ChiS$ R1030Q, Tm<sup>R</sup> |
| SAD 3026 | Figure 2 GFP R1046Q | $\Delta lacZ::P_{chb}$-GFP, Kan<sup>R</sup>; igVCA0265/0266::CBS-mCherry v2, Cm<sup>R</sup>; $\Delta chiS/cbp::Carb^R$; $\Delta VCA0692::ChiS$ R1046Q, Tm<sup>R</sup> |
| SAD 3027 | Figure 2 GFP K1052Q | $\Delta lacZ::P_{chb}$-GFP, Kan<sup>R</sup>; igVCA0265/0266::CBS-mCherry v2, Cm<sup>R</sup>; $\Delta chiS/cbp::Carb^R$; $\Delta VCA0692::ChiS$ K1052Q, Tm<sup>R</sup> |
| SAD 3028 | Figure 2 GFP R1063Q | $\Delta lacZ::P_{chb}$-GFP, Kan<sup>R</sup>; igVCA0265/0266::CBS-mCherry v2, Cm<sup>R</sup>; $\Delta chiS/cbp::Carb^R$; $\Delta VCA0692::ChiS$ R1063Q, Tm<sup>R</sup> |
| SAD 3029 | Figure 2 GFP R1068Q | $\Delta lacZ::P_{chb}$-GFP, Kan<sup>R</sup>; igVCA0265/0266::CBS-mCherry v2, Cm<sup>R</sup>; $\Delta chiS/cbp::Carb^R$; $\Delta VCA0692::ChiS$ R1068Q, Tm<sup>R</sup> |
| SAD 3030 | Figure 2 GFP R1074Q | $\Delta lacZ::P_{chb}$-GFP, Kan<sup>R</sup>; igVCA0265/0266::CBS-mCherry v2, Cm<sup>R</sup>; $\Delta chiS/cbp::Carb^R$; $\Delta VCA0692::ChiS$ R1074Q, Tm<sup>R</sup> |
| SAD 3031 | Figure 2 GFP K1078Q | $\Delta lacZ::P_{chb}$-GFP, Kan<sup>R</sup>; igVCA0265/0266::CBS-mCherry v2, Cm<sup>R</sup>; $\Delta chiS/cbp::Carb^R$; $\Delta VCA0692::ChiS$ K1078Q, Tm<sup>R</sup> |
| SAD 3032 | Figure 2 GFP R1081Q | $\Delta lacZ::P_{chb}$-GFP, Kan<sup>R</sup>; igVCA0265/0266::CBS-mCherry v2, Cm<sup>R</sup>; $\Delta chiS/cbp::Carb^R$; $\Delta VCA0692::ChiS$ R1081Q, Tm<sup>R</sup> |
| SAD 3033 | Figure 2 GFP K1087Q | $\Delta lacZ::P_{chb}$-GFP, Kan<sup>R</sup>; igVCA0265/0266::CBS-mCherry v2, Cm<sup>R</sup>; $\Delta chiS/cbp::Carb^R$; $\Delta VCA0692::ChiS$ K1087Q, Tm<sup>R</sup> |
| SAD 3034 | Figure 2 GFP R1090Q | $\Delta lacZ::P_{chb}$-GFP, Kan<sup>R</sup>; igVCA0265/0266::CBS-mCherry v2, Cm<sup>R</sup>; $\Delta chiS/cbp::Carb^R$; $\Delta VCA0692::ChiS$ R1090Q, Tm<sup>R</sup> |
| SAD 3035 | Figure 2 GFP R1092Q | $\Delta lacZ::P_{chb}$-GFP, Kan<sup>R</sup>; igVCA0265/0266::CBS-mCherry v2, Cm<sup>R</sup>; $\Delta chiS/cbp::Carb^R$; $\Delta VCA0692::ChiS$ R1092Q, Tm<sup>R</sup> |
| SAD 3036 | Figure 2 GFP K1106Q | $\Delta lacZ::P_{chb}$-GFP, Kan<sup>R</sup>; igVCA0265/0266::CBS-mCherry v2, Cm<sup>R</sup>; $\Delta chiS/cbp::Carb^R$; $\Delta VCA0692::ChiS$ K1106Q, Tm<sup>R</sup> |
| SAD 3037 | Figure 2 GFP R1112Q | $\Delta lacZ::P_{chb}$-GFP, Kan<sup>R</sup>; igVCA0265/0266::CBS-mCherry v2, Cm<sup>R</sup>; $\Delta chiS/cbp::Carb^R$; $\Delta VCA0692::ChiS$ R1112Q, Tm<sup>R</sup> |
| SAD 3038 | Figure 2 GFP R1119Q | $\Delta lacZ::P_{chb}$-GFP, Kan<sup>R</sup>; igVCA0265/0266::CBS-mCherry v2, Cm<sup>R</sup>; $\Delta chiS/cbp::Carb^R$; $\Delta VCA0692::ChiS$ R1119Q, Tm<sup>R</sup> |
| SAD 3039 | Figure 2 GFP K1121Q | $\Delta lacZ::P_{chb}$-GFP, Kan<sup>R</sup>; igVCA0265/0266::CBS-mCherry v2, Cm<sup>R</sup>; $\Delta chiS/cbp::Carb^R$; $\Delta VCA0692::ChiS$ K1121Q, Tm<sup>R</sup> |
| SAD 3040 | Figure 2 GFP K1124Q | $\Delta lacZ::P_{chb}$-GFP, Kan<sup>R</sup>; igVCA0265/0266::CBS-mCherry v2, Cm<sup>R</sup>; $\Delta chiS/cbp::Carb^R$; $\Delta VCA0692::ChiS$ K1124Q, Tm<sup>R</sup> |
| SAD 2675 | Figure S2 ChiS<sup>WT</sup> No FLAG tag | $\Delta lacZ::P_{chb}$-GFP, Kan<sup>R</sup>; $\Delta chiS/cbp::Carb^R$; $\Delta VCA0692::ChiS$, Tm<sup>R</sup> |

**Table S4.**
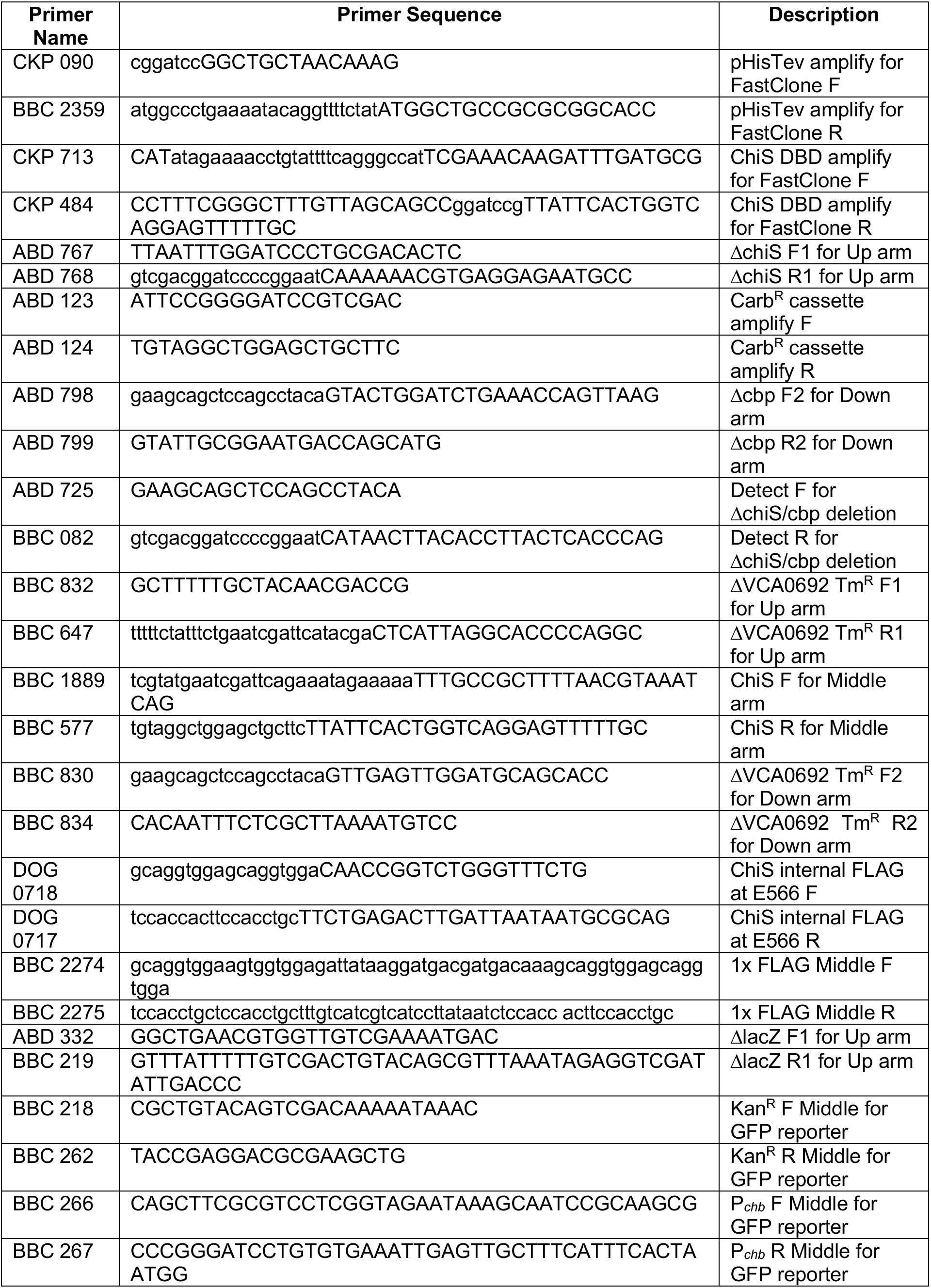
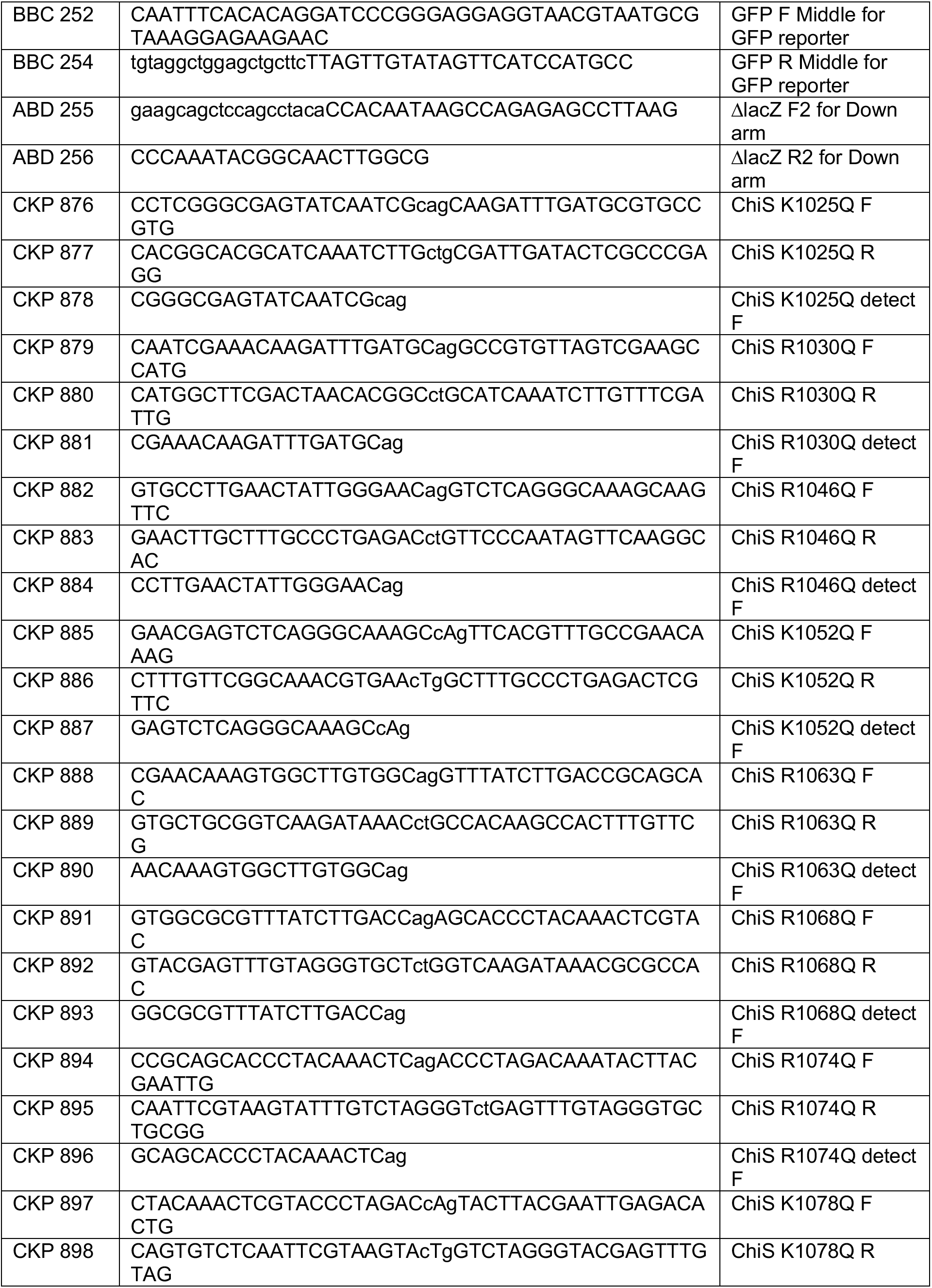
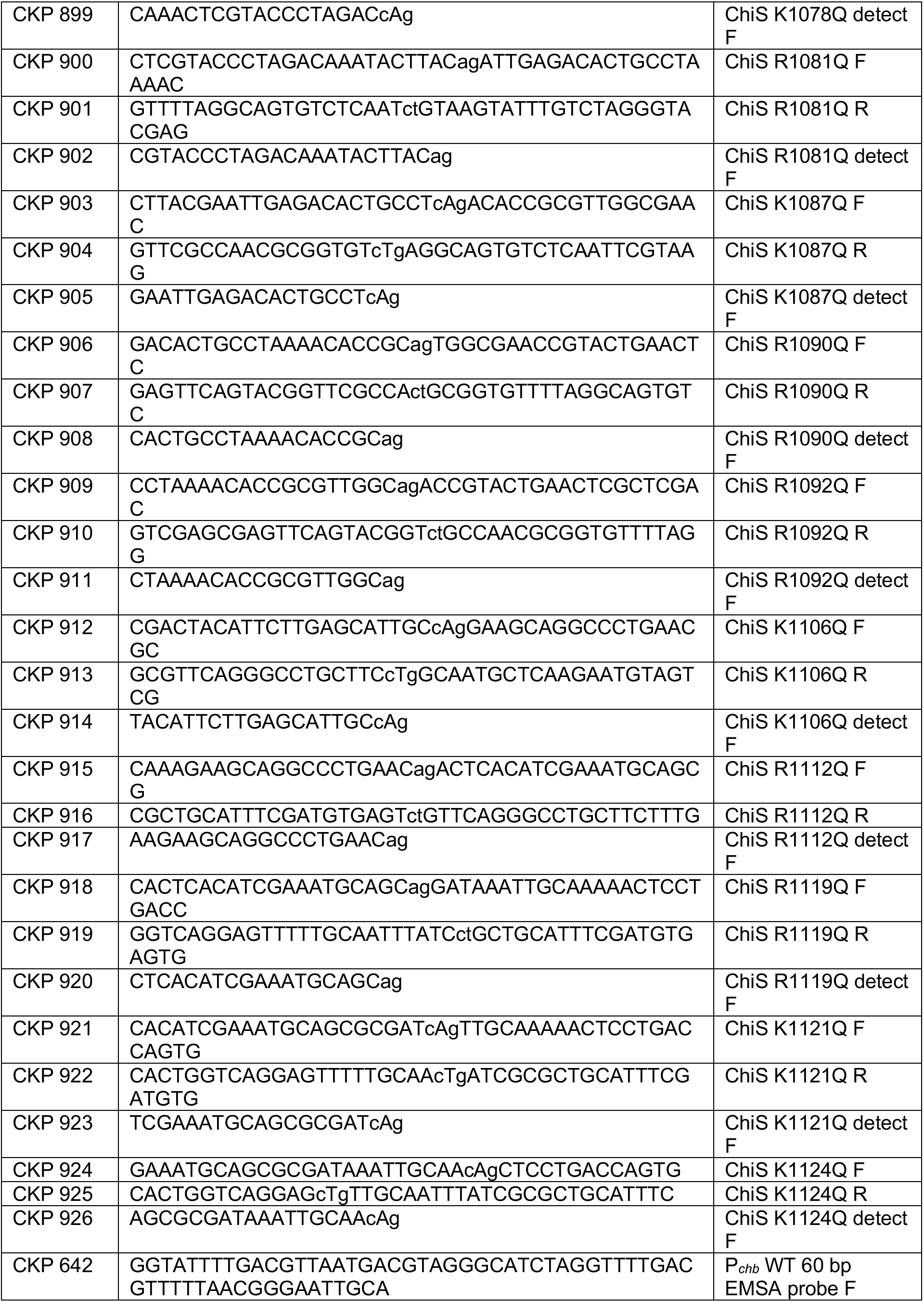
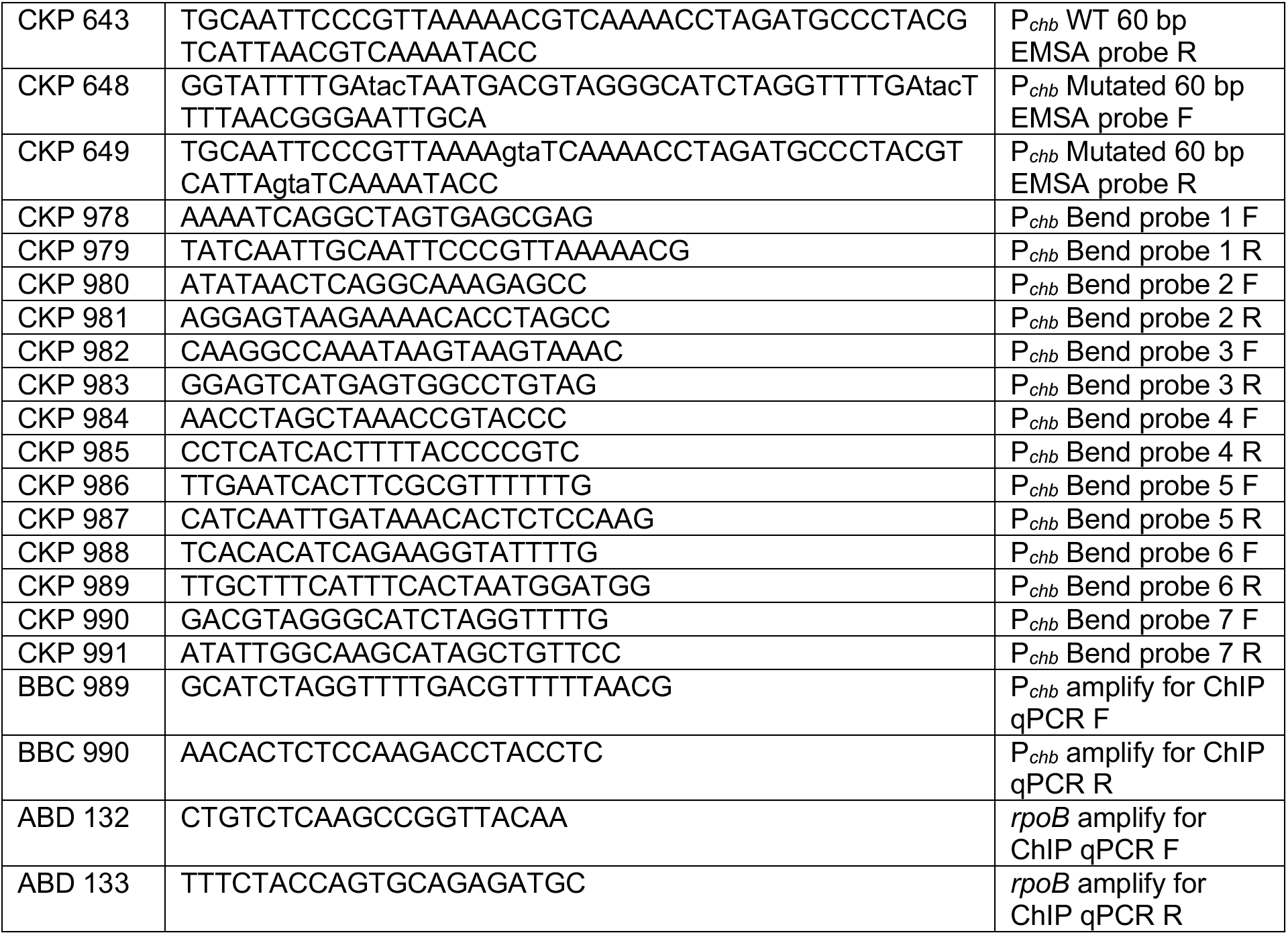
Primers used in this study.

